# Protein network analysis links the NSL complex to Parkinson’s disease and mitochondrial biology

**DOI:** 10.1101/2023.01.27.524249

**Authors:** Katie Kelly, Patrick A. Lewis, Helene Plun-Favreau, Claudia Manzoni

## Abstract

Whilst the majority of PD cases are sporadic, much of our understanding of the pathophysiological basis of disease can be traced back to the study of rare, monogenic forms of disease. In the past decade, the availability of Genome-Wide Association Studies (GWAS) has facilitated a shift in focus, toward identifying common risk variants conferring increased risk of developing PD across the population. A recent mitophagy screening assay of GWAS candidates has functionally implicated the non-specific lethal (NSL) complex in the regulation of PINK1-mitophagy. Here, a bioinformatics approach has been taken to investigate the proteome of the NSL complex, to unpick its relevance to PD progression. The mitochondrial NSL interactome has been built, mining 3 separate repositories: PINOT, HIPPIE and MIST, for curated, literature-derived protein-protein interaction (PPI) data. We built; i) the ‘mitochondrial’ interactome, applying gene-set enrichment analysis (GSEA) to explore the relevance of the NSL mitochondrial interactome to PD and, ii) the PD-oriented interactome to uncover biological pathways underpinning the NSL /PD association. In this study, we find the mitochondrial NSL interactome to be significantly enriched for the protein products of PD associated genes, including the Mendelian PD genes *LRRK2* and *VPS35*. Additionally, the PD associated interactome is enriched for mitochondrial processes; *“mitochondrial cell death*”, *“mitochondrial protein localisation”*, “*membrane protein localisation”* and *“mitochondrial transport”*. Our data points to NSL complex members OGT and WDR5 as key drivers of this increased PD association. These findings strengthen a role for mitochondrial quality control in both familial and sporadic disease.

## Introduction

Parkinson’s disease (PD) is the most common movement disorder of old age (> 65 years) ^1^. Further, global prevalence predictions suggest that the number of affected individuals more than doubled in 25 years, with an estimated 6.1 million people living with PD in 2016 ^2^.

The movement aspects of PD are triggered by the progressive degeneration of neurons within the Substantia Nigra pars compacta (SNpc). The consequent depletion of Dopamine (DA) within the nigro-striatal circuits gives origin to the debilitating triad of PD clinical symptoms: rigidity, asymmetric resting tremor, and bradykinesia. Pathologically, neuronal loss is paired with the deposition of α-synuclein amyloid aggregates in intracellular inclusions called Lewy bodies ^3^. The progression of PD is complex, with involvement of additional brain areas whose degeneration is responsible for the clinical manifestation of additional non-motor symptoms ^4^. PD has a heterogenous presentation and the interindividual differences in disease onset and progression, typical of complex disorders, hint at the existence of a personal burden of risk, a mix of genetic susceptibility factors and environmental exposures which differ on a case-to-case basis ^4^. Potentiation of DA signalling is the only available therapeutic intervention, achieved via DA replacement and/or by inhibition of DA catabolism and re-uptake at the synapse. However, this is a symptomatic intervention that does not halt neurodegeneration ^5^. As such, precise delineation of the molecular mechanisms of PD neurodegeneration is critical, to implicate novel research avenues for development of disease-modifying treatments.

A minority of PD cases are familial (fPD), caused by highly penetrant mutations that segregate with the disease. The study of monogenic forms of PD has facilitated the elucidation of common cellular phenotypes, and the molecular patterns underpinning them. Functional delineation of the *PINK1* and *PRKN* genes, for which loss-of-function mutations are causal for autosomal recessive (AR) PD, has implicated dysfunctional mitophagy as one of the key drivers of disease ^6, 7^. PINK1 and Parkin act in concert to promote the targeted degradation of depolarised mitochondria ^8-11^. Additionally, mutations within the AR PD gene *DJ-1*, as well as parkinsonism genes *FBXO7* and *VPS35*, can also be functionally associated with mitochondrial quality control ^12^. While *LRRK2* and *GBA (*associated with late onset forms of PD) have been linked to dysregulation of autophagy and the endolysosmal pathway ^13, 14^. A comprehensive understanding of the relationship between genes and phenotypes in fPD has implicated biological processes that could be relevant for disease. However, as monogenic forms of PD represent approximately 5-10% of all cases ^3^, the question arises as to how appropriately we can model PD in the absence of a comprehensive understanding of the molecular processes underpinning sporadic disease (sPD).

To this end, data from genome-wide association studies (GWAS) provide an unbiased approach to investigate common genetic variation and disease risk across the population. To-date, the identification of > 80 independent genetic risk signals has supported the existence of a genetic architecture, conferring an increased personal risk of developing sPD ^15^. However, GWAS pinpoint risk loci, rather than specific genes, so the molecular pathways behind this genetic architecture remain elusive. Translation of genetic data to the precise delineation of relevant molecular pathways represents a rate limiting step in studies of this kind.

Recently, our group has identified, using a High-Content Screening assay, a functional association between KAT8 (otherwise known as MOF), a MYST family Lysine acetyltransferase, and PINK1-mitophagy ^16^. The *KAT8* gene is located at the 16q 11.2 PD risk locus, near one of the genome wide significant PD risk signals. KAT8 is one of nine components (HCFC1, KANSL1, KANSL2, KANSL3, KAT8, MCRS1, OGT, PHF20, WDR5) of the Non-Specific Lethal (NSL) complex. While a nuclear role for the NSL complex has been well defined, a study by Chatterjee et al suggests partial localisation of the complex to the mitochondria ^17^. Indeed, Knock Down (KD) of NSL members KANSL1, KANSL2, KANSL3 and MCRS1 have also been shown to impede PINK1-dependent induction of mitophagy ^16^. Interestingly, the *KANSL1* gene is an additional PD-GWAS hit ^18^, located at the inversion polymorphism on chromosome 17q21, alongside *MAPT*. Up to 25% of individuals of European descent inherit, within this region, a sequence of ∼1mb, in the opposite orientation ^19, 20^. This induces a ∼1.3–1.6 Mb region of linkage disequilibrium (LD), preventing recombination. Thus, haplotype-specific polymorphisms have resulted in the emergence of two major haplotype clades, H1 (the most common) and H2, of which H1 has been strongly linked to neurodegenerative disease, including PD ^21-23^. While *MAPT*, encoding the tau protein, is frequently attributed to the PD risk association at this locus ^24^, evidence has accrued supporting a contribution of *KANSL1* to risk at this locus. In summary, there is strong evidence that functional alterations of members of the NSL complex might underpin the risk signal for sPD at these two loci. These findings provide functional evidence for the importance of mitophagy and mitochondrial quality control in sporadic forms of disease, in addition to fPD. To gain a greater insight into the functional links between the NSL complex, mitophagy and PD, we have constructed an *in-silico* protein-protein interaction (PPI) model of the NSL complex, describing its relationship with the mitochondrial proteome in the context of the PD genetic landscape.

Our analysis reveals that the intersection between the mitochondrial proteome and the NSL interactome is enriched with PD genes. Additionally, the interactome centred around PD genes in direct connection with the NSL complex, is enriched for mitochondrial processes. We therefore propose that the alteration in the functionality of the NSL complex at the mitochondria is the one of the molecular causes of sPD. Therefore, alteration in mitophagy can be seen as molecular bridge between familial and sporadic forms of disease.

## Materials and Methods

The methods for the *in silico* analysis of the NSL complex in Parkinson’s disease and mitochondria are detailed in protocols.io: dx.doi.org/10.17504/protocols.io.5qpvorb19v4o/v1

### Nomenclature

A layered approach has been taken to build the NSL protein network (*NSL-PN*), whereby members of the NSL complex are designated as seeds to derive a list of proteins for which a physical interaction has been experimentally determined. To achieve this, interactors of NSL complex members, termed Protein-Protein Interactors (PPIs), have been downloaded, filtered, and prioritised.

Herein, the NSL complex members are designated as *‘NSL seeds’. NSL seeds* along with the *first layer* interactors, will be termed the *‘first layer’*. The *‘second layer’* constitutes direct interactors of *first layer* members, proteins that are linked to the *NSL seeds* via a “*bridge*”, which is a protein in the *first layer*. The “*interactome*” of an individual seed comprises the seed under investigation, with the direct interactors (in the *first layer*), plus the indirect interactors (in the *second layer*).

*NSL seeds*, plus the *first* and *second* layers, comprise the complete *‘NSL-PN’*. Within this analysis, we have taken two approaches to generate the complete *NSL-PN*; generating a *‘Mito-CORE network*’, and a ‘*PD-CORE network*’ (outlined below). PD associated *first layer* members will be termed ‘*PD seeds’* and mitochondrial *first layer* members, *‘Mito seeds’*. Within this text, we refer to a complete list of 19,947 genes (obtained by processing the file downloaded from HGNC; https://www.genenames.org/download/statistics-and-files/ [August 2019]) as the ‘*Gene dictionary’* ^25^. The pipeline for building each layer of the network has been described hereafter.

### Downloading the protein-protein interaction (PPI) data

The pipeline to derive the *first layer* interactome is displayed in Figure 1. PPIs for *NSL seeds* (Supplementary table 1; DOI: <u>10.5281/zenodo.7516686</u>) were collected using 3 different web-based tools; PINOT v1.1 (Protein Interaction Network Online Tool; https://www.reading.ac.uk/bioinf/PINOT/PINOT_form.html [downloaded September 2021 using the lenient filter option])^25^, HIPPIE with no threshold on interaction score (Human Integrated Protein-Protein Interaction rEference; RRID:SCR_014651; http://cbdm-01.zdv.uni-mainz.de/∼mschaefer/hippie/network.php; DOI: 10.1371/journal.pone.0031826 [downloaded September 2021]) ^26^ and MIST v5.0 (Molecular Interaction Search Tool; https://fgr.hms.harvard.edu/MIST [downloaded September 2021]) ^27^. Each resource permitted interrogation of a selection of IMEx consortium associated repositories, to obtain literature-derived, curated PPI data.

**Figure 1.**
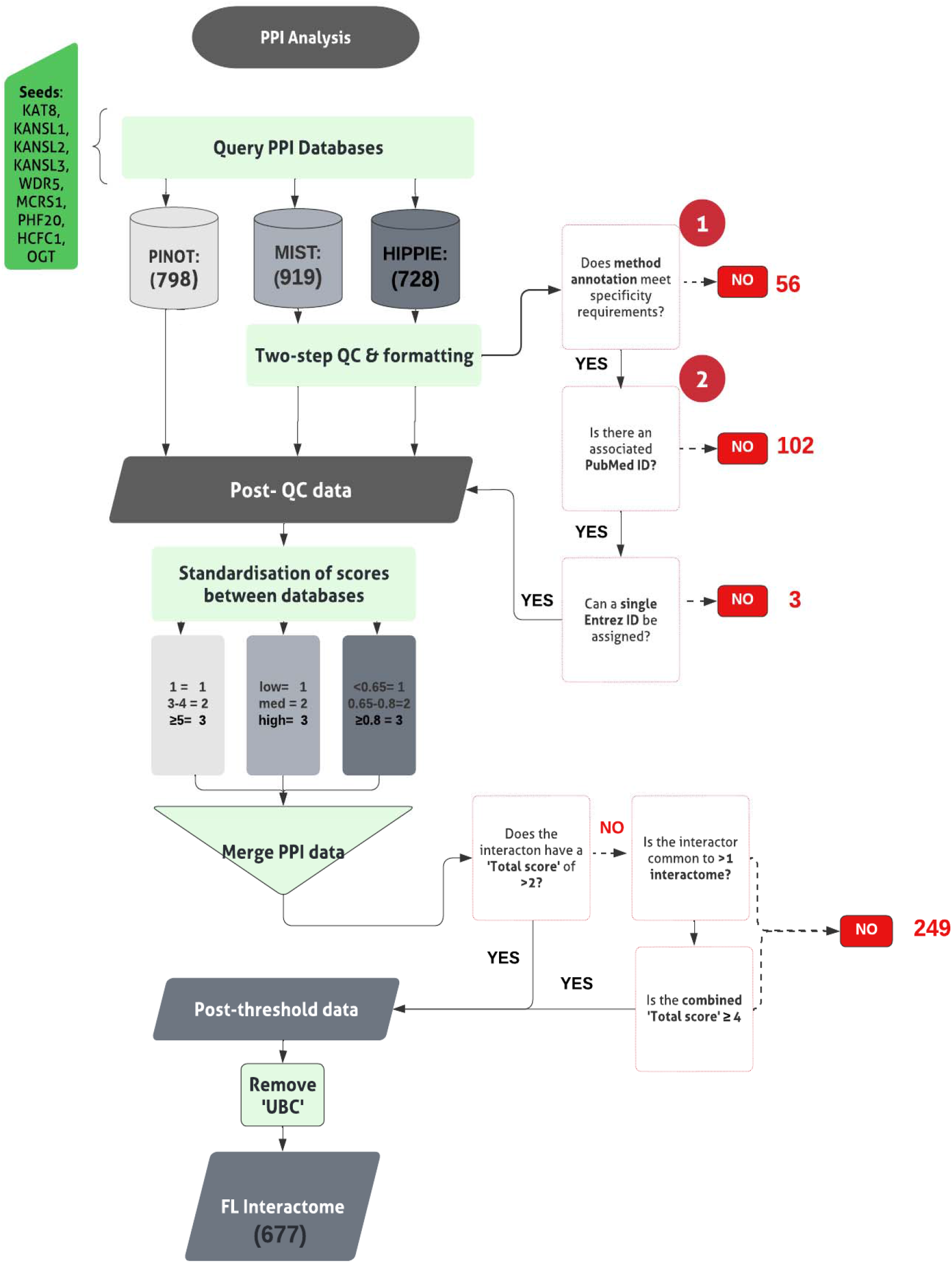
Pipeline for building the *first layer* interactome of the NSL complex. The ‘Seeds’ are the nine members of the NSL complex. Circled numbers (1 & 2) indicate the two stages of quality control (QC) applied. Numbers in black font, provided in brackets, indicate total number of interactions retained at each stage. Numbers red font indicate the number of interactions removed at each stage.

PPI data obtained using MIST and HIPPIE has been subjected to quality control (QC), already integrated within the PINOT pipeline, to remove low quality data. Entries lacking *“interaction detection method”* annotation (QC1), or a PubMed ID (QC2) have been removed. Formatting between the output files has been standardized and interactors’ IDs converted to the approved EntrezID, UniprotID and HGNC gene name using the *Gene dictionary*. Proteins with nonunivocal conversions to these 3 identifiers were removed.

Where ‘UBC’, a ubiquitin moiety, was identified as an interactor within the *first layer*, we manually reviewed the supporting publication. Ubiquitin is understood to be conjugated to proteins as a marker for degradation. As such, these were considered as potentially introducing non-specific protein interactions into the analysis. UBC was removed from the *first layer* interactomes of OGT and WDR5, since both interactions have been identified via high throughput, as opposed to specific, methods.

### Merging and thresholding the PPIs

Prior to merging the lists of PPIs obtained from the different databases, a confidence score (CS) was assigned to each of the interactions. The continuous CS of HIPPIE and PINOT were converted into the categorical CS of MIST. Low CS was then assigned to a value of 1, moderate CS was assigned to a value of 2 and, high CS was assigned to a value of 3.

Interaction data, across the 3 databases, was merged to generate a single file for each seed’s interactome. For each interaction the (*CS*_*T*_) was calculated as:

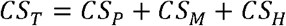

The (*CS*_*T*_) ranges therefore from a min = 1 (PPI reported in only 1 database with low confidence), to a max = 9 (PPI reported by the 3 databases, always with high confidence).

An arbitrary score threshold (*CS*_*T*_ >2) was then applied, to filter and remove lower confidence PPI data lacking reproducibility. For an interaction to have a *CS*_*T*_ = 3, it must be reported either with low confidence across all 3 databases, or with moderate or high confidence in a single database. In the case that the interaction is reported with low confidence across 3, we reason that it has at a minimum passed the stringent QC of PINOT, and thus have retained the interaction.

Interactions that failed to meet the threshold were interrogated further, to identify those interactors bridging >1 interactome. For those interactors appearing within >1 interactome, a multi-interactome threshold represented by a *CS*_*T*_ ≥ 4 across interactomes, was applied. Those meeting this multi-interactome threshold were retained.

### Generating the Mito-CORE network

The pipeline for steps taken to derive the ***Mito-CORE network*** can be found in Figure 2. First, we prioritised members of the *first layer* with mitochondrial annotation (- OGT, since it was a seed to derive the *first layer* interactome), which we termed ‘*Mito seeds’*. Proteins with mitochondrial annotation were obtained via 2 independent inventories:

**Figure 2.**
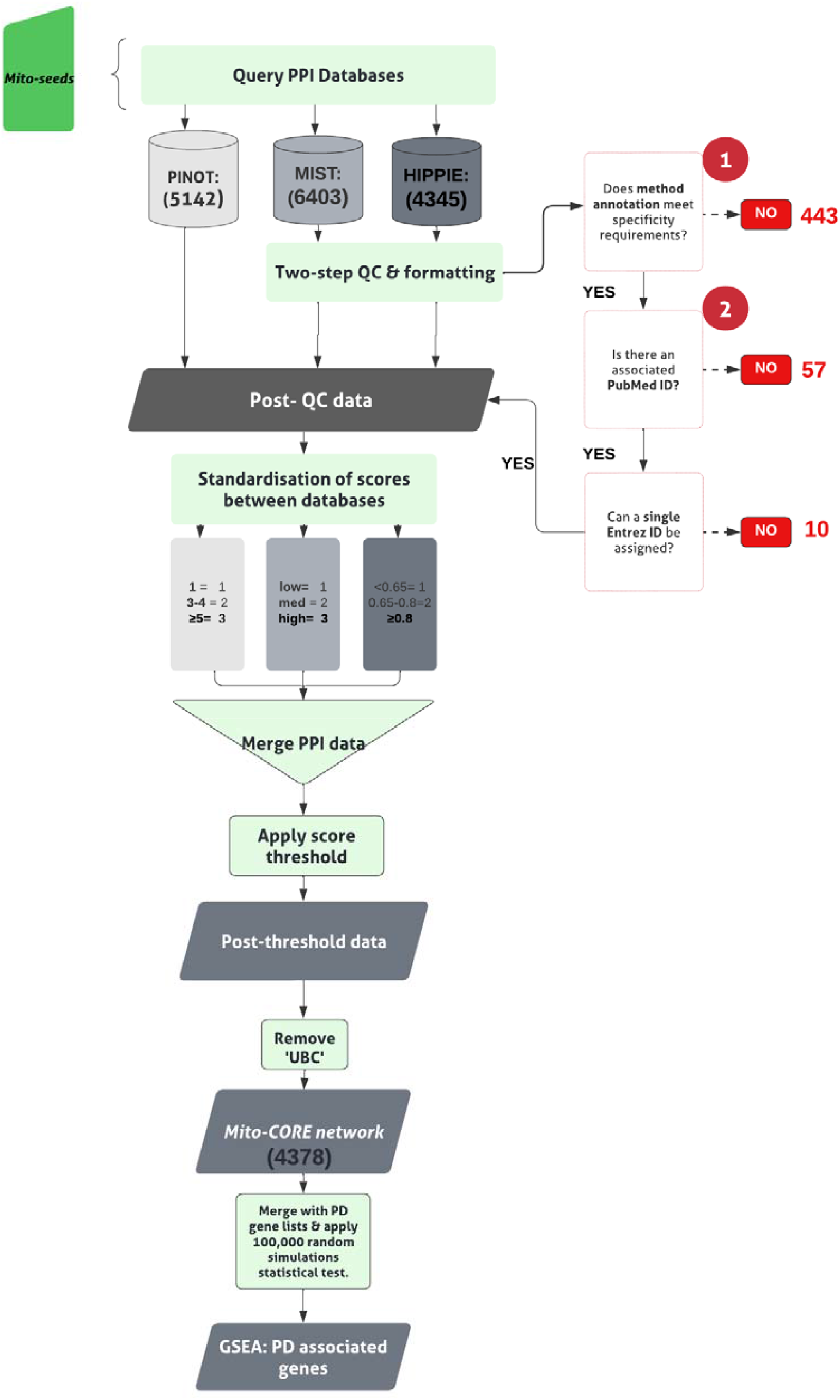
Pipeline for building the *Mito-CORE*, and application of PD Gene-set enrichment analysis (GSEA). *‘Mito-seeds’* refers to the mitochondrial *first layer* members of the NSL interactome. Circled numbers (1 & 2) indicate the two stages of quality control (QC) applied. Numbers in black font, provided in brackets, indicate total number of interactions retained at each stage. Numbers red font indicate the number of interactions removed at each stage. * Score threshold is applied as described in the pipeline in Figure 1, after the ‘Merge PPI data’ step.

i)the AmiGO2 encyclopedia ^28, 29^ (RRID:SCR_002143) was queried (February 2022), to derive experimentally determined mitochondrial protein lists. Two accession terms were used: GO: 0005759, to obtain proteins annotated to the “mitochondrial matrix” and GO:0031966 for proteins annotated to the “mitochondrial membrane”; in both cases, ‘Homo sapiens’ was specified as the search organism. ii) the Human MitoCarta3.0 dataset ^30^ (RRID:SCR_018165) was downloaded (October 2021) to retrieve proteins for which a Mitochondrial Targeting Sequence (MTS) has been identified. Interactors’ IDs were converted to the approved EntrezID, UniprotID and HGNC gene name using the *Gene dictionary*. Proteins with nonunivocal conversions to these 3 identifiers were removed.

*Mito seeds* were input into all three PPI tools, to obtain the *second layer* (November 2021). The *NSL seeds* together with the *Mito seeds*, and *second layer* interactors formed the complete ***Mito-CORE network***.

### Gene Set Enrichment Analysis (GSEA)

GSEA for PD associated genes was conducted, by comparing the members of the interactome under investigation to a list of 180 unique PD associated genes; generated by consulting 3 publicly accessible resources: i) PanelApp v1.68 diagnostic grade genes (green annotations) for PD and Complex Parkinsonism ^31^ (Gene Panel: Parkinson’s Disease and Complex Parkinsonism (Version 1.108); https://panelapp.genomicsengland.co.uk/panels/39/ [downloaded October 2021]) and ii) the latest GWAS meta-analysis ^15^ as well as iii) a list of 15 genes associated with Mendelian PD, obtained from a recent W-PPI-NA ^32^. The genes from i, ii, and iii were combined, to generate a PD associated genes list, herein referred to as the ‘PD genes’ list (Supplementary table 2 ; DOI: <u>10.5281/zenodo.7516686</u>). The genes within this list have been referred to by the name of their protein product.

### Statistical evaluation via random networks simulation

To test the significance of gene set overlap, 100,000 random simulations were carried out and used to validate statistical significance of overlaps of PD genes with the *first layer* and complete *Mito-CORE network*. 100,000 random gene lists, each of them equivalent in length to *first layer*/complete *Mito-CORE network*, were obtained using the R random sampling function, from the *Gene dictionary*. Each list was compared to the PD gene list, keeping track of the matches. The *p*-value has been calculated using the *p-*norm function in R, which compares the real number of experimental matches to the actual number of experimental matches.

### Generating the PD-CORE network

The pipeline is reported in Figure 3. The genes in the intersection between the *first layer* and the list of 180 unique PD genes were used as *PD seeds*. They were input into PINOT to obtain the *second layer* interactome. An arbitrary confidence threshold of ‘*CS*_p_ >2’ has been applied, eliminating data with just a single publication and method from the downstream analysis. Once again interactors’ IDs were converted to the approved EntrezID, UniprotID and HGNC gene name using the *Gene dictionary*. Proteins with nonunivocal conversions to these 3 identifiers were removed. To remove background noise, only members of the *second layer* bridging >1 *PD seed* within the *PD-CORE network* were kept. Protein interactors that were private to 1 *PD seed* only were removed, this resulted in the removal of WDR5B from the *PD-CORE network*. The *NSL seeds* together with the *PD seeds*, and the non-private *second layer* interactors have thus formed the complete ***PD-CORE network***.

**Figure 3.**
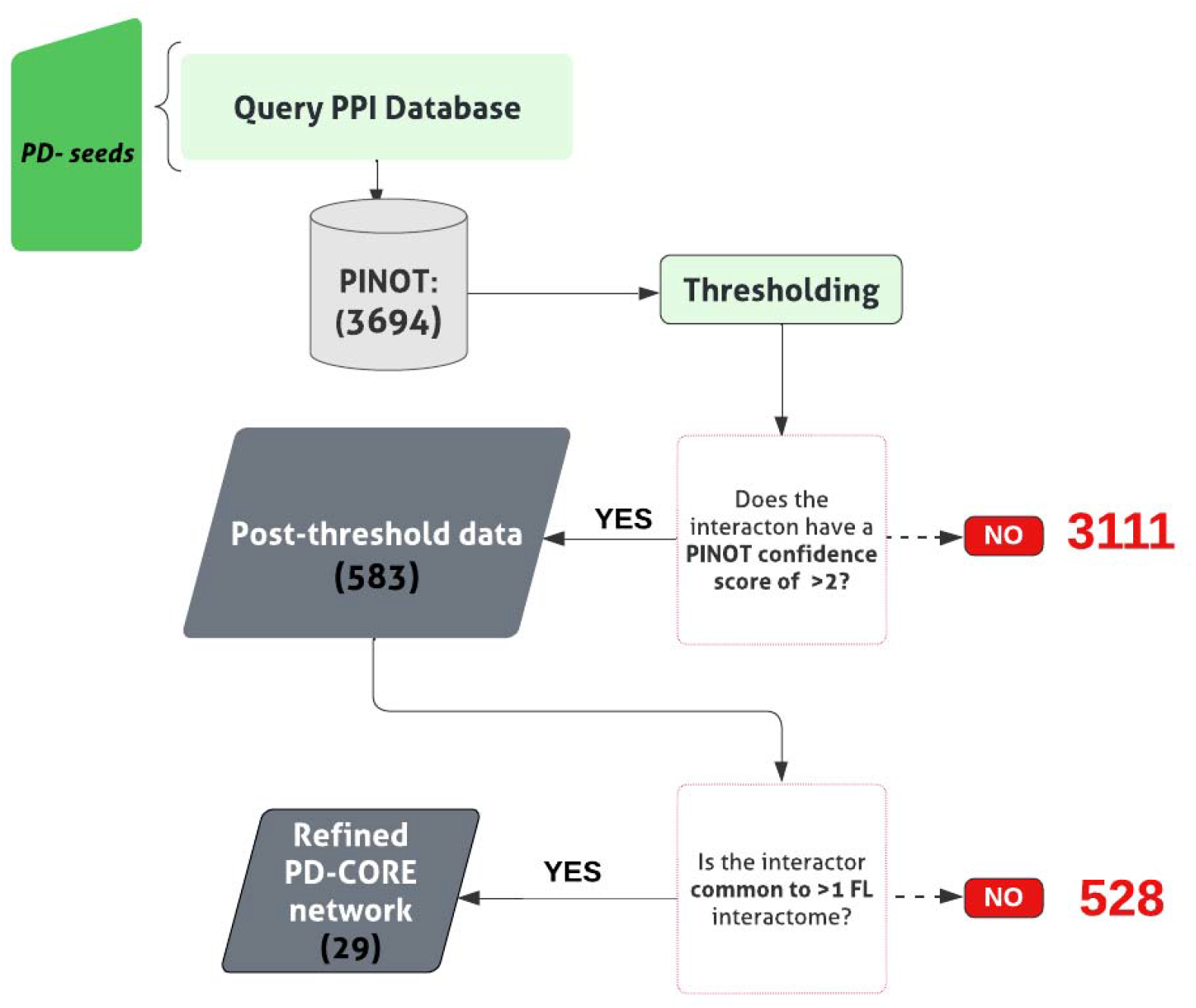
Pipeline for building the *PD-CORE*. *‘PD-seeds’* refers to the PD associated *first layer* members. Numbers in black font, provided in brackets, indicate total number of interactions retained at each stage. Numbers red font indicate the number of interactions removed at each stage.

### Functional enrichment analysis

To assess enrichment of particular biological processes within the *PD-CORE network*, members *(- NSL seeds*), were input into the g:Profiler search tool ^28, 29^ (g:GOSt; RRID:SCR_006809 http://biit.cs.ut.ee/gprofiler/ [downloaded January 2022]). Enrichment for GO terms associated with ‘Biological Processes (BPs)’ only, was conducted, with all other analysis settings left unadjusted, generating a list of enriched GO:BP terms.

A threshold was applied to the list of enriched GO:BP terms, to retain those with term size <100 thus effectively removing ‘broad’ GO:BP terms. Remaining terms were assigned to custom-made ‘semantic classes’(SC), accompanied by a parent ‘functional group’(FG). Generic terms (classified in the semantic classes of: General, Metabolism, and response to Stimulus) were discarded from further analysis. The pipeline for this analysis is presented in Supplementary figure 1. GO:BP terms contributing to each SC were pooled, to identify the list of proteins within the network contributing to the enrichment of that specific SC.

### Software and scripts

Where data has been parsed in Rstudio (version 1.3.1093; *RRID:SCR_000432;* http://www.rstudio.com/) the script can be obtained at: https://doi.org/10.5281/zenodo.7346957 within the relevant project files. Additional software used are Microsoft Excel (version 16.6; *RRID:SCR_016137;* https://www.microsoft.com/en-gb/) and Cytoscape (version 3.8.2; *RRID:SCR_003032;* http://cytoscape.org/) ^33^.

## Results

### Construction of the NSL-PN: First Layer

To construct the *first layer* of the NSL protein network (*NSL-PN*), the nine members of the NSL complex served as seeds to query three separate PPI repositories: PINOT, MIST and HIPPIE, obtaining a set of direct interactors of the NSL complex. Three tools were consulted at this stage, to maximize capture of PPI data available within the literature. Search from PINOT, MIST and HIPPIE yielded a total of 798, 919 and 728 direct interactions, respectively. To pool the data between the three search tools, differences in formatting and protein identification nomenclature needed to be standardized. Each Interactor Identifier (ID) was converted to the approved EntrezID, approved UniprotID and HGNC gene name; proteins with nonunivocal conversions to these 3 IDs were removed. To standardise the quality of data between tools, it was necessary to apply two quality control (QC) steps to data captured by MIST and HIPPIE excluding data without a publication ID and/or an associated interaction detection method. A summary of excluded and retained data can be found in Table 1. Once data collected from PINOT, MIST and HIPPIE was pooled, 948 unique interactions were observed across the 9 NSL interactomes (Supplementary table 1), of which 63% were captured by all three search tools (Supplementary figure 2). Each single seed-interactome contained the following number of interactors: KAT8: 45, KANSL1: 56, KANSL2: 23, KANSL3:18, PHF20: 23, WDR5:256, MCRS1: 184, OGT: 181, HCFC1: 162.

**Table (1).** *NSL seeds* with corresponding UniProt ID; used to interrogate three separate protein-protein interaction (PPI) tools (PINOT,MIST & HIPPIE). Columns contain PPI counts; column headings correspond to stage of the analysis. QC1 = Quality Control step 1; step to remove data lacking ‘method annotation’. QC2 = Quality Control step 2; step to remove data lacking PubMed ID. Here, column ‘-UBC’ corresponds to total PPI count after removal of this ubiquitin moiety from the final interactor list.

| NSL Seed | UniProt ID | PPIs downloaded (P,H,M) | Removed : QC1 | Removed : QC2 | Removed: un-resolved multi-entrez ID | Unique PPIs [post QC & formatting] | Post-threshold PPIs | -UBC |
| --- | --- | --- | --- | --- | --- | --- | --- | --- |
| KAT8 | Q9H7Z6 | 38,42,49 | 5 | 2 | 1 | 45 | 41 | 41 |
| KANSL1 | Q7Z3B3 | 49,48,54 | 2 | 0 | 0 | 56 | 48 | 48 |
| KANSL2 | Q9H9L4 | 20,13,22 | 1 | 0 | 0 | 23 | 18 | 18 |
| KANSL3 | Q9P2N6 | 13,13,18 | 1 | 0 | 0 | 18 | 17 | 17 |
| PHF20 | Q9BVI0 | 16,16,23 | 3 | 0 | 1 | 23 | 19 | 19 |
| WDR5 | P61964 | 230,226,249 | 18 | 9 | 1 | 256 | 215 | 214 |
| MCRS1 | Q96EZ8 | 177,153,175 | 0 | 89 | 0 | 184 | 88 | 88 |
| OGT | O15294 | 140,153,175 | 14 | 2 | 0 | 181 | 138 | 137 |
| HCFC1 | P51610 | 115,134,152 | 12 | 0 | 0 | 162 | 116 | 116 |

Steps were then taken to refine each interactome, to remove ‘low quality’ data. Each of the databases have scoring systems in place to evaluate the ‘confidence’ of each interaction. The confidence score (CS) represents an evaluation of the amount of evidence (e.g., number of publications/methods reporting the interaction, type of method used to evaluate the interaction) supporting a given interaction. To obtain a ‘Total score (*CS*_*T*_)’ for each interaction, it was necessary to elaborate the individual CS assigned by PINOT (*CS*_*p*_), MIST (*CS*_*M*_) and HIPPIE (*CS*_*H*_ ). Since each database uses a different scoring system; we opted to convert the continuous scores assigned by PINOT and HIPPIE to the categorical scores of MIST (1=low, 2 = moderate 3 = high) prior to calculation of the *CS*_*T*_ (please refer to materials and methods), which ranged from a min= 1, to a max = 9. To remove lower confidence interactions, an arbitrary score threshold of ‘*CS*_*T*_ > 2’ was applied. An interaction with a score of 1 or 2 indicates that it has either been reported with low confidence in two out of three repositories, or that it has been reported with moderate confidence in just one; in either case, the interaction was considered as lacking support by literature evidence. A concessionary threshold was applied to those interactors that did not meet this threshold but were common to more than one seed interactome (a combined threshold of ‘*CS*_*T*_ > 4’ across interactomes was used in this case). This provided a means to prioritise interactors associating with more than one NSL complex member; interactions that could be important for delineating a function of the NSL complex, rather than its constituent members. Following refinement of the data, ∼74% of interactions were retained across all interactomes (Supplementary figure 3).

To the post threshold total, each single seed interactome contributed the following number of interactors: KAT8: 41, KANSL1: 48, KANSL2: 18, KANSL3:17, PHF20: 19, WDR5:214, MCRS1: 88, OGT: 137, HCFC1: 116. Merging of the nine interactomes generated the *first layer* of the *NSL-PN*; represented by 517 single nodes (i.e., unique interactors), and 677 undirected edges (i.e., unique interactions). None of the nine interactomes within the *first layer* were isolated, all were connected in the same unique graph, supporting the functional association of all seeds as part of the NSL complex.

### The first layer interactome is enriched for PD associated genes

We sought to assess whether there was an enrichment of proteins associated with PD in the *first layer*; by overlapping the 517 *first layer* members, with the list of 180 PD genes (Supplementary table 2, see methods for details). Indeed, an intersection of 13 proteins between the *first layer* and the list of PD genes was found (*p*-value = 4.45 × 10^−5^). The intersection remained significant even when calculated with a more stringent list containing only the 15 Mendelian PD genes (see methods for details) (*p*-value= 4.5 × 10^−3^) (Supplementary table 3). Out of the 13 PD genes represented within the *first layer*, the protein products of *LRRK2* and *VPS35*, mutations in which cause autosomal dominant (AD) forms of PD, were found to be direct interactors of the NSL complex. Taken together, these findings strengthen the argument for a functional association between the NSL complex and the disease mechanisms that underpin PD.

### Construction of the NSL-PN: Second Layer

To minimise bias deriving from the use of NSL complex members as seeds (i.e., seed centrality bias), a multi-layered *NSL-PN* was built (i.e., *first layer* plus *second layer* interactors). Two different approaches were taken to prioritise members of the *first layer*, and to generate an expansion of the network to the *second layer* of protein interactions. First, a ‘mitochondrial’ NSL interactome was built, referred to as the ***‘Mito-CORE network’***, to explore the relevance of the NSL mitochondrial interactome to PD. Secondly, a ‘PD-oriented’ NSL interactome was built, referred to as the ***‘PD-CORE network’***, to uncover biological pathways associated with the portion of the NSL complex network that is relevant for PD.

### The Mito-CORE network is enriched for PD associated genes

To establish the *Mito-CORE network*, members of the *first layer* interactome were prioritised based on mitochondrial annotation; the pipeline for building this network can be followed in Figure 2. A list of 1346 unique ‘mitochondrial proteins’ were derived from two independent inventories; the AmiGO2 encyclopaedia and the Human MitoCarta3.0 dataset (Supplementary table 4). Overlapping this list of 1364 proteins with the components of the *first layer* NSL interactome revealed an intersection of 22 proteins (Figure 4A). A list of 21 mitochondrial proteins within the *first layer* were used as *‘Mito-seeds’* and represented the starting point to download the *second layer* interactors using PINOT, MIST and HIPPIE (upon exclusion of OGT) (a summary of downloaded data can be found in Table 2). The resultant *Mito-CORE network* held 2832 single nodes (unique interactors) and 4373 undirected edges (unique interactions) (Supplementary table 5; DOI: 10.5281/zenodo.7516686). Interestingly, there was a modest increase in the mean connections for each single node (n of edges/n of nodes) from 1.30 in the first layer, to 1.54 in the *Mito-CORE network*. We also observed an increase in the average number of neighbours with the addition of the *second layer*, from 2.6 to 3.1, indicating an increase in functional connectedness in the *Mito-CORE network*. We next sought to assess whether an enrichment of genes associated with PD was upheld in the complete *Mito-CORE network*. Notably, there were 45 overlaps between the PD gene list, and the complete *Mito-CORE network*, accounting for 25 % of the complete PD gene list being represented (Figure 4B). Thus, there is significant enrichment of proteins encoded by PD associated genes within the mitochondrial interactome of the NSL complex (*p-*value= 1.48 × 10^−5^; Supplementary figure 4) (complete list of overlaps reported in Supplementary table 6). Moreover, 8/15 genes from the more ‘stringent’ Mendelian PD gene list were represented in the *Mito-CORE network*: *GBA, PRKN, SNCA, PRKRA, PARK7, FBXO7, VPS35 and LRRK2* (Figure 4B), an overlap which meets statistical significance (*p-*value = 6.81 × 10^−6^; Supplementary figure 5). Considering these results, we propose that the mitochondrial interactome of the NSL complex might represent one of the functional links between the NSL complex and PD.

**Table (2).** *Mito seeds* with corresponding UniProt ID; used to interrogate three separate PPI tools (PINOT,MIST & HIPPIE), to generate the *second layer* of the complete *Mito-CORE network*. Columns contain data counts; column headings indicate the corresponding stage of the analysis. QC1 = Quality Control step 1; step to remove data lacking ‘method annotation’. QC2 = Quality Control step 1; step to remove data lacking PubMed ID. Here column ‘-UBC’ corresponds to total PPI count after removal of this ubiquitin moiety from the final interactor list.

| Mito Seed | UniProt ID | PPIs downloaded (P,H,M) | Removed: QC1 | Removed: QC2 | Removed: un-resolved multi-entrez ID | Unique PPIs [post QC & formatting] | Post-threshold PPIs |
| --- | --- | --- | --- | --- | --- | --- | --- |
| HSPD1 | P63038 | 649,665,726 | 254 | 2 | 2 | 767 | 639 |
| MRPS15 | P82914 | 70,66,77 | 1 | 0 | 0 | 82 | 75 |
| MRPL11 | Q9Y3B7 | 95,83,106 | 30 | 16 | 1 | 110 | 70 |
| FOXRED1 | Q96CU9 | 28,26,51 | 0 | 0 | 0 | 52 | 34 |
| MAVS | Q7Z434 | 108,121,142 | 23 | 0 | 0 | 150 | 112 |
| SNAP29 | O95721 | 95,91,108 | 11 | 8 | 0 | 112 | 93 |
| ECI2 | O75521 | 81,89,100 | 8 | 1 | 0 | 100 | 82 |
| LAP3 | P28838 | 46,40,80 | 7 | 9 | 0 | 84 | 40 |
| DUS2 | Q9NX74 | 4,6,7 | 0 | 0 | 0 | 7 | 4 |
| **AGMAT** | Q9BSE5 | 22,24,27 | 9 | 0 | 0 | 27 | 20 |
| **FOXO3** | O43524 | 71,80,100 | 2 | 0 | 0 | 100 | 84 |
| **LRRK2** | Q5S007 | 1439,548,1850 | 5 | 1 | 1 | 1861 | 638 |
| **PLEC** | Q15149 | 238,238,278 | 16 | 1 | 1 | 290 | 248 |
| **PPP1CC** | P36873 | 345,371,651 | 5 | 3 | 2 | 660 | 383 |
| **PPP2R2B** | Q00005 | 191,198,201 | 2 | 0 | 1 | 206 | 198 |
| **RPS27A** | P62979 | 178,200,223 | 51 | 12 | 0 | 263 | 172 |
| **TRAK1** | Q9UPV9 | 17,22,21 | 3 | 0 | 0 | 23 | 20 |
| **PPP1R15A** | O75807 | 128,130,162 | 1 | 0 | 0 | 167 | 131 |
| **TERT** | O14746 | 73,108,114 | 1 | 1 | 0 | 119 | 104 |
| **TP53** | Q96S44 | 1101,1073,1190 | 15 | 3 | 2 | 1316 | 1064 |
| **CCAR2** | Q8N163 | 163,166,189 | 10 | 0 | 0 | 205 | 167 |

**Figure 4.**
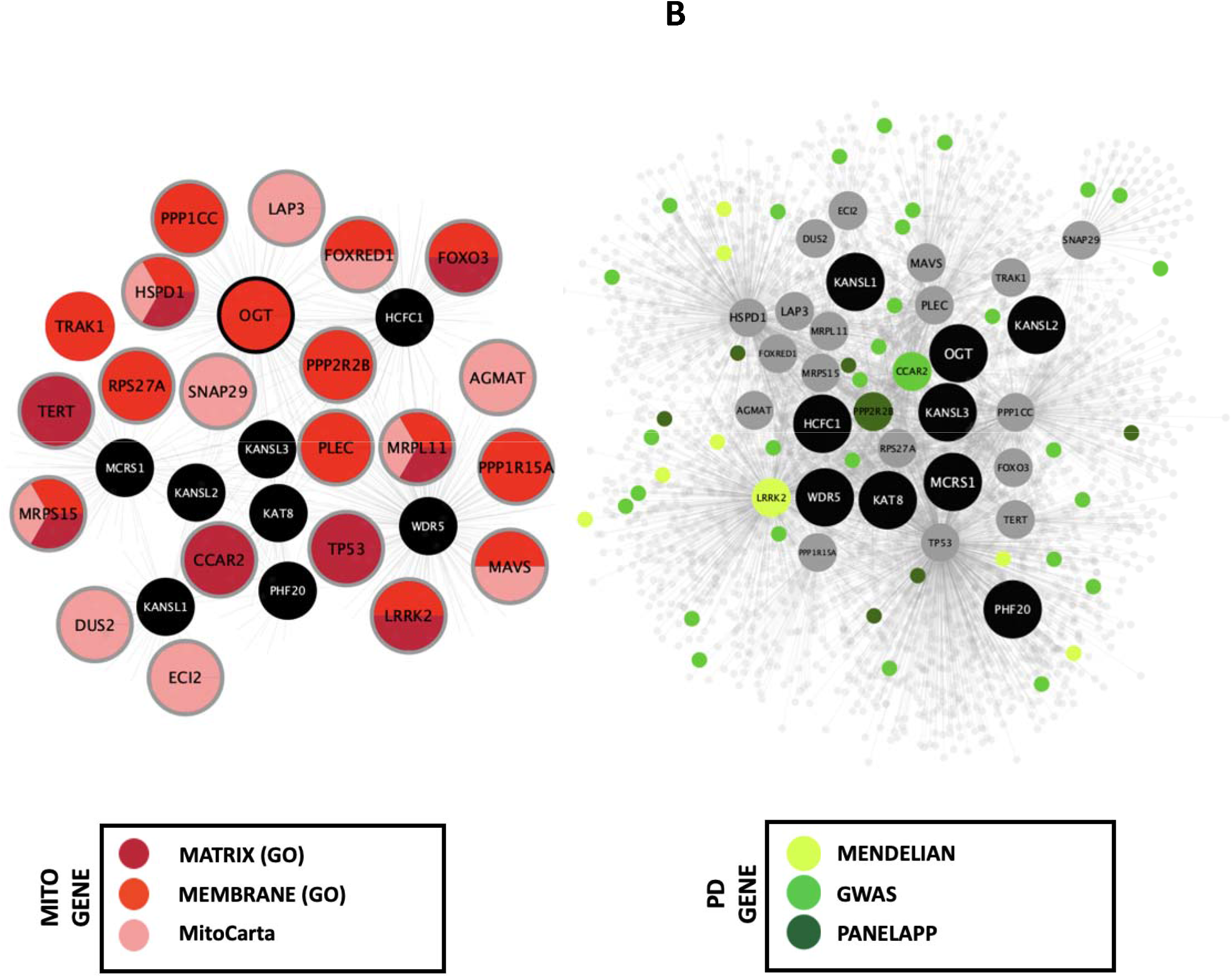
The *Mito-CORE network* is enriched for PD-associated gene products. *(A)* 21 members of the *first layer* interactome were prioritised as *‘Mito seeds*’ for derivation of the *second layer*. Nodes are colour coded according to the repository reporting mitochondrial localisation **(B)** The *Mito-CORE network* was subject to PD gene set enrichment analysis (GSEA), to reveal statistically significant enrichment of PD risk genes (45/180; 25%) (*p-*value= 1.48 × 10^−5^). We also found 8 out of a stringent list of 15 genes associated with Mendelian PD, represented within the *Mito-CORE network:* GBA, PRKN, SNCA, PRKRA, PARK7, FBXO7, VPS35 and LRRK2, an enrichment which meets statistical significance (*p* value = 6.81 × 10^−6^).

### The PD-CORE network is enriched for mitochondrial processes

We next expanded the *PD-CORE* (*NSL seeds*, with the 13 PD associated *first layer* members) to generate the ***PD-CORE network***. The pipeline for building this network is illustrated in Figure 3. Input of the PD associated *first layer* members into PINOT followed by QC (a summary of retained data can be found in Table 3) initially returned a total of 583 protein interactions (Supplementary table 7; DOI: 10.5281/zenodo.7516686). At this stage, an additional refinement step was applied to remove *second layer* nodes connected to <2 *first layer* interactors; thus, keeping those proteins in the *second layer* that were able to bind multiple PD associated proteins of the *first layer* (Supplementary table 8; DOI: 10.5281/zenodo.7516686). This step increased the connectivity of the network, removing *second layer* members which might represent ‘background noise’. The *PD-CORE network* contained 71 nodes comprising the *NSL seeds*, 12/13 of the PD associated *first layer* members, and their interactions in the *second layer* (communal to more than 2 PD proteins in the *first layer*) (Figure 5).

**Table (3).** *PD Seeds* with corresponding UniProt ID; used to interrogate PINOT to generate the second layer of the *PD-CORE network*. ‘PD association’ column, indicates the source of the association. ‘Mendelian’: refers to a list of 15 genes associated with Mendelian PD. GWAS: refers to a list of PD risk genes from the latest PD GWAS meta-analysis. ‘PANELAPP’: refers to a list of diagnostic grade genes for PD and Complex Parkinsonism. Columns contain data counts; column headings indicate the corresponding stage of the analysis. ‘Post-threshold PPIs’ correspond to those with PINOT confidence scores >2. ‘PD-association threshold’ corresponds to the number of PPIs bridging >1 interactome within the *PD-CORE network*. WDR5B was removed from *the PD-CORE network*, once private interactors were excluded.

| **PD Seed** | **UniProt ID** | **PD association**<br>**(G=GWAS, P=PanelApp,**<br>**W=W PPI NA)** | **Total PPIs**<br>**downloaded** | **Post-**<br>**threshold**<br>**PPIs** | **Post PD-**<br>**association**<br>**threshold** |
| --- | --- | --- | --- | --- | --- |
| **CAMK2D** | Q13557 | G | 153 | 17 | 3 |
| **CSTA** | P01040 | G | 66 | 6 | 1 |
| **CYLD** | Q9NQC7 | G | 612 | 32 | 7 |
| **ATN1** | P54259 | P | 104 | 15 | 4 |
| **MAPT** | P10636 | G/P | 620 | 84 | 34 |
| **PPP2R2B** | Q00005 | P | 191 | 85 | 15 |
| **UBTF** | P17480 | G | 87 | 19 | 2 |
| **SETD1A** | O15047 | G | 37 | 15 | 4 |
| **WDR5B** | Q86VZ2 | G | 26 | 1 | 0 |
| **VPS35** | Q96QK1 | P/W | 91 | 18 | 1 |
| **CCAR2** | Q8N163 | G | 163 | 26 | 6 |
| **SGF29** | Q96ES7 | G | 105 | 53 | 3 |
| **LRRK2** | Q5S007 | G/P/W | 1439 | 212 | 40 |

**Figure 5.**
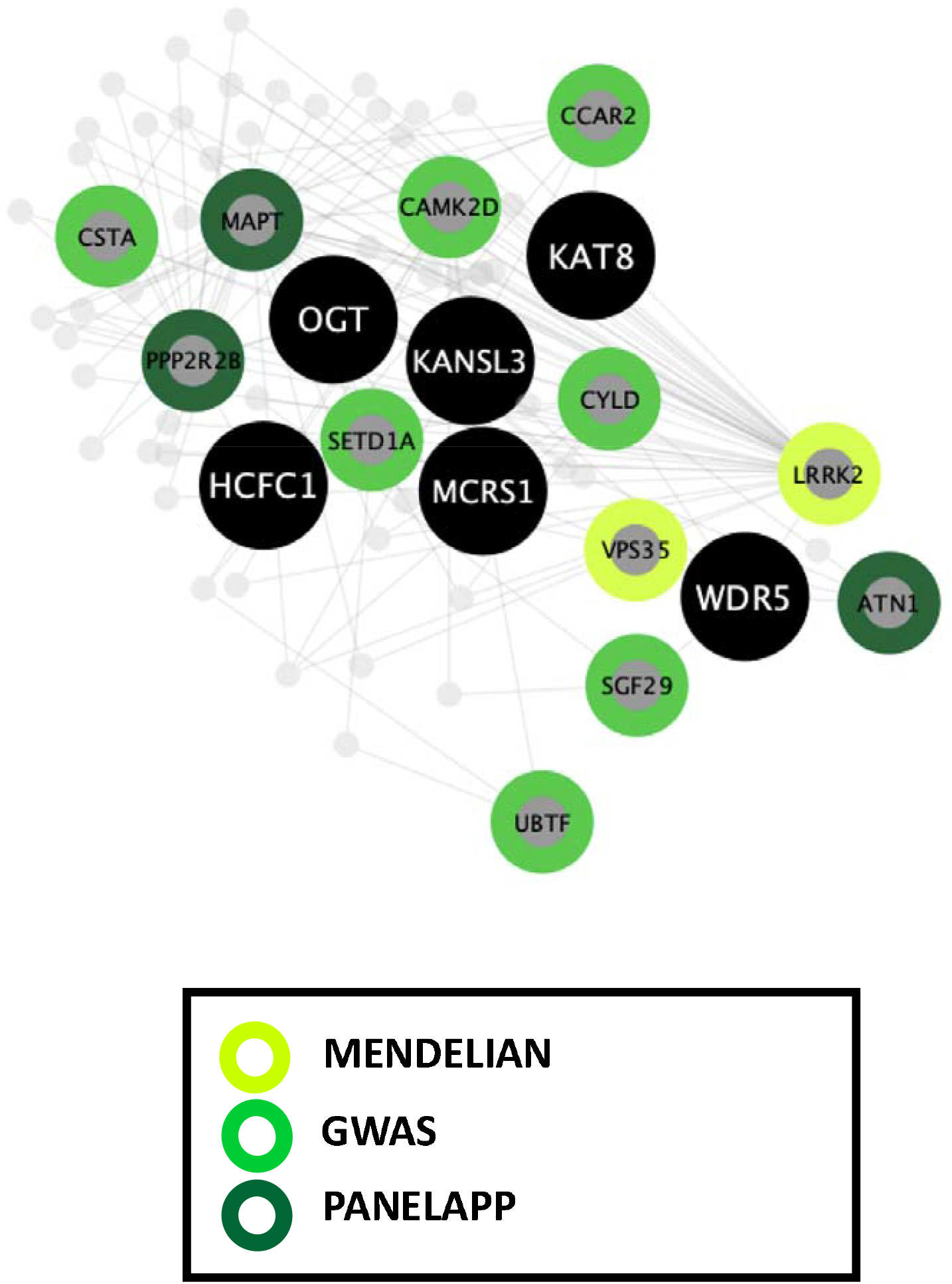
Pipeline for building the *PD-CORE network*. The *PD-CORE network* was derived by prioritising the 13 PD-associated members of the NSL-PN *first layer*. The *second layer* was obtained, to generate the *PD-CORE network*. Subsequent refinement of the network, removing ‘private’ interactors, generated the refined *network*, comprised of 12 *first layer* interactors (*PD-seed* WDR5B was removed) and 53 *second layer* interactors of seeds; KAT8, KANSL3, OGT, WDR5, HCFC1, MCRS1 (black nodes). ‘Mendelian’: refers to a list of 15 genes associated with Mendelian PD. GWAS: refers to a list of PD risk genes from the latest PD GWAS meta-analysis. ‘PANELAPP’: refers to a list of diagnostic grade genes for PD and Complex Parkinsonism.

To determine enriched biological processes within the *PD-CORE network*, its 65 members (excluding the *NSL seeds*) were input into the g: Profiler search tool (g:GOSt). 405 GO: BP terms were returned, 45 of which were retained after subsequent refinement to remove broader terms (term size >100). Once each GO: BP had been manually assigned to a ‘functional group’ (FG) and ‘semantic class’ (SC) enrichment of 9 FGs and 20 SCs was observed (Supplementary table 9) (Figure 6A and B). The most significantly enriched SC (in terms of *p*-value and total number of single GO: BPs allocated to them) were *“mitochondrial cell death*”, *“mitochondrial protein localisation”*, “*membrane protein localisation”* and *“mitochondrial transport”*. It is notable that 3 of the 4 most enriched processes, with the highest weight and statistical significance, pertain to mitochondrial processes (Figure 6B). Taken together, this analysis reveals that mitochondrial processes are strongly associated with the proteins composing the *PD-CORE network*. The *PD-CORE network* was then refined, retaining only those (29 + 4 *NSL seeds*) proteins responsible for this enrichment (Figure 7A), allowing extraction of the PD - mitochondrial subnetwork. Topological analysis of the extracted subnetwork showed that two of the *NSL seeds* (OGT and WDR5) contributed to much of the network with 83% (24/29)) and 93% (27/29) of the *PD-CORE* nodes represented by the OGT and WDR5 interactomes respectively, while only 21% (6/29) and 7% (2/29) of the *PD-CORE* nodes are represented by the HCFC1 and KAT8 interactomes (Figure 7B). Taken together, these findings point to OGT and WDR5 as key drivers of the PD association with the NSL mitochondrial processes, at the protein level.

**Figure 6.**
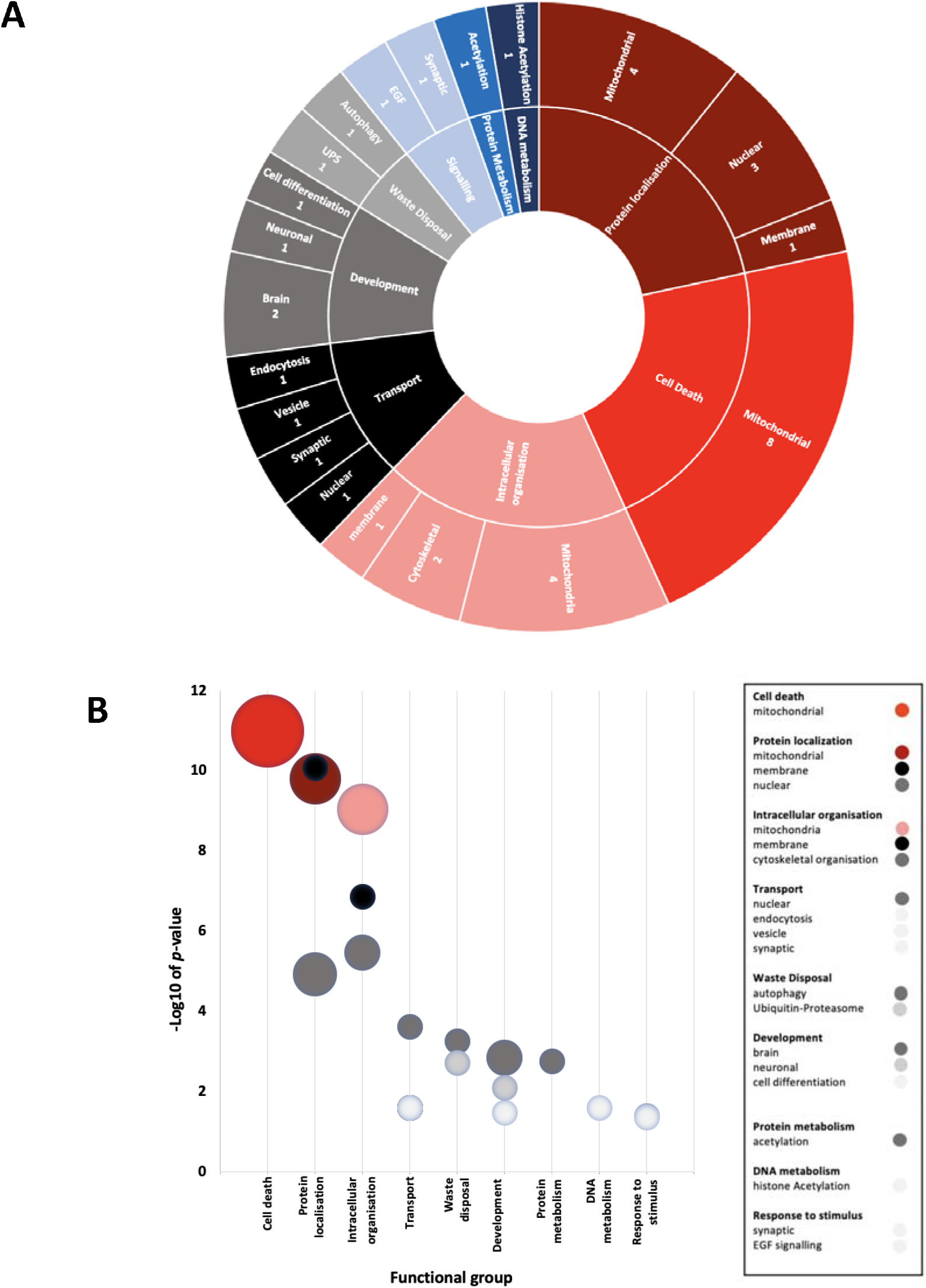
Mitochondrial processes are functionally enriched within the *PD-CORE network*. **(A)** The pie chart depicts the proportion of enriched GO terms represented by each Functional Group (FG) (outside ring), and each Semantic Class (SC), inner ring. FGs for which there is mitochondrial representation amongst SCs are depicted in a shade of red. Numbers correspond to the number of GO terms represented by each SC **(B)** The bubble chart illustrates weighted SCs. The lowest *p*-value of associated GO terms has been allocated to the SC. Bubble size represents the number of GO terms within the SC.

**Figure 7.**
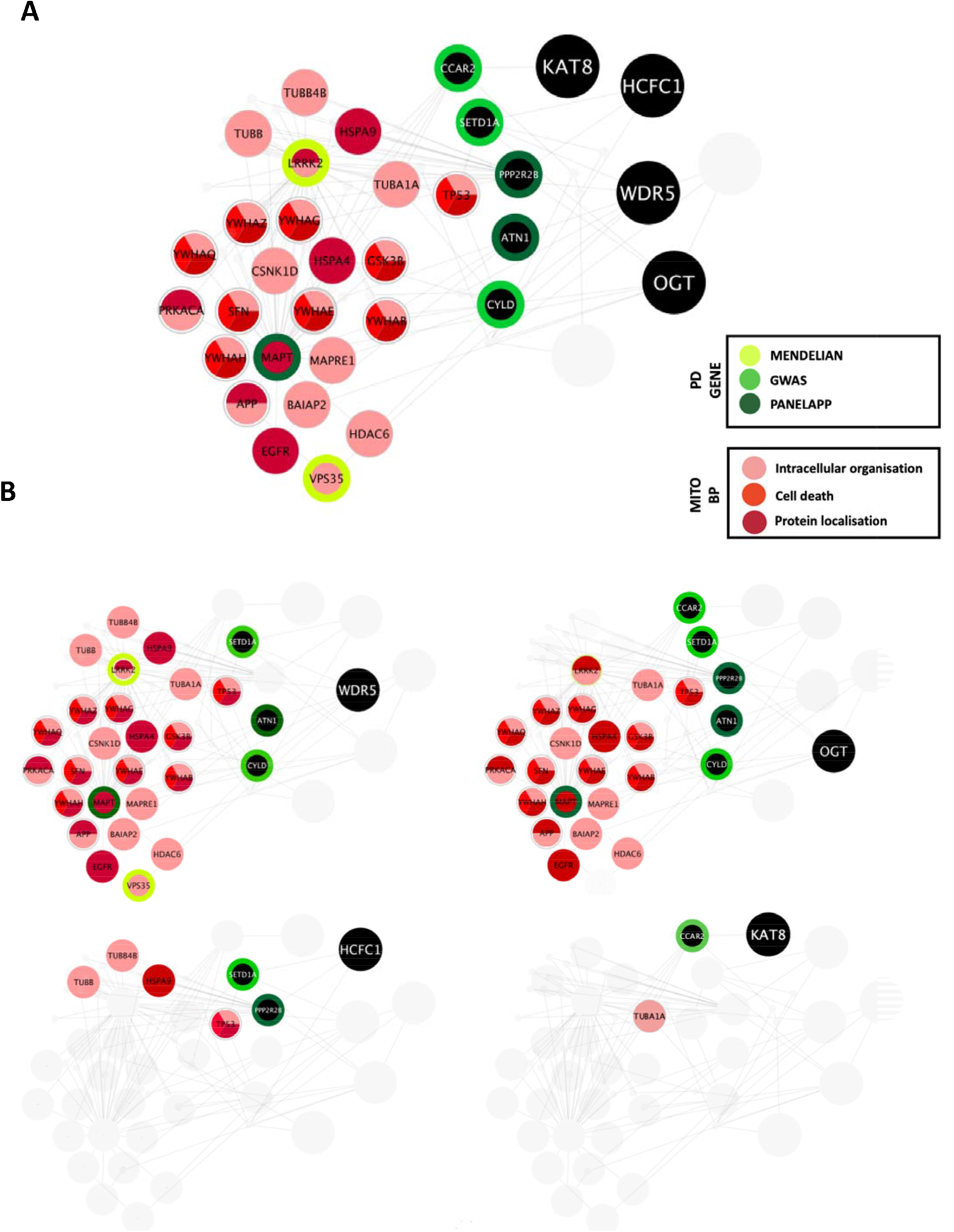
OGT and WDR5 are possible key drivers of the PD risk associated with mitochondrial processes. **(A)** Visualisation of the refined of the *PD-CORE network*, containing proteins that contribute to the enrichment of mitochondrial processes. Final network contains 29 nodes (FL + SL) + 4 NSL seeds. ‘MITO BP’ refers to ‘mitochondrial biological process’ **(B)** extraction and isolation of contributing interactomes, demonstrates that OGT and WDR5 contribute most significantly to the enrichment of mitochondrial processes with 83% (24/29)) and 93% (27/29) of the *PD-CORE* nodes represented by the OGT and WDR5 interactomes, respectively.

## Discussion

To date, functional studies of the genetic basis for PD have focused on delineating the precise molecular underpinnings of monogenic PD. However, the genetic architecture of both familial and sporadic PD is complex, and an interplay between genetic and environmental factors has been proposed to be the basis of PD aetiopathogenesis ^4^. While GWAS have revealed multiple genetic risk factors, these studies identify loci as opposed to specific genes, which are associated with an increase or decrease in the risk of PD development. Recently, our lab has revealed a role for several members of the NSL complex in the regulation of PINK1-mitophagy; for which 2 members have been proposed as genetic risk factors for sporadic PD development (Soutar et al. 2022) ^15^. Thus, it has been suggested that the NSL complex could play a role in mitochondrial quality control mechanisms within the context of sporadic PD. To gain a deeper understanding of how the NSL complex intersects with pathways involved in PD, in the present study we re-constructed the protein interactome of the NSL complex. Our results demonstrate multiple functional links to mitochondrial dynamics, as well as other PD associated genes, providing a computational prediction that the NSL complex, via mitochondrial pathways, serves as a link between familial and sporadic PD.

Protein-protein interactions (PPIs) were collected from peer-reviewed literature to construct the *NSL-PN*. Initially, we used three separate databases to obtain a comprehensive list of direct interactors for each of our nine *NSL seeds*, to construct the *first layer* interactome. PPI data has utility in guiding the interpretation of data generated by genomics studies, to make biological sense of the data, inferring functional associations and pathways associated with disease risk. However, PPI studies are inherently limited by ascertainment bias, since data available reflects interactions that have been determined by hypothesis-led experimentation within the wet lab, resulting in an over-representation of interactions reported for proteins with an existing disease-association ^34^. Indeed, in the present study, we report 215 interactors for WDR5, while only 17 for KANSL3 and 18 for KANSL2. The disparity in interactome size that we have found could reflect the physiologically relevant difference in the number of cellular interactors of each protein, however it is not possible to exclude the possibility that it might reflect bias in the research literature. To minimise the effect of this research bias within our analysis, we have taken a multi-layered approach to build the *NSL-PN*. Rather than looking at individual members of the interactome, we have carried out a set of analyses using the entire *NSL-PN*, suitable for drawing meaningful conclusions from an inherently partial dataset.

In a first approach, we expanded upon the *first layer* of the *NSL-PN* to obtain the ***Mito-CORE network*** of the NSL complex. The NSL complex, with KAT8, is well characterized as a master regulator of transcription, responsible for acetylation of histone 4 at lysine 5, lysine 8 and lysine 16 at the nuclear compartment ^35, 36^. However, a role for the NSL complex at the mitochondria has also been suggested ^17^. This suggestion is of interest in the context of PD, since mitochondrial quality control/dynamics have been intimately associated with familial PD ^37^. We therefore filtered the *NSL-PN* to retain only mitochondrial proteins. The identification of mitochondrial proteins was conducted using two independent inventories that were pooled to maximise coverage. We retrieved the Human MitoCarta3.0 dataset ^38^, a set of proteins harbouring a mitochondrial targeting sequence (MTS), along with candidates obtained from the AmiGO2 encyclopaedia ^28, 29^, experimentally evidenced to localise to the mitochondrial matrix or membrane. Using this approach, we found 21/517 members of the *NSL-PN first layer* to be localised to the mitochondria; ECI2, DUS2, PPP1CCC, PPP2R2B, MAVS, LRRK2, TP53, SNAP29, RPS27A, MRPL11, MRPS15, AGMAT, FOXO3, FOXRED1, LAP3, HSPD1, TRAK1, TERT, CCAR2, PPP1R15A and OGT. While 21/517 represents a lower proportion than would be expected by chance, we suggest that the incomplete nature of PPI data alongside a research bias toward a nuclear function for the NSL complex could explain this lack of data. We used these mitochondria localised proteins within the *first layer* of the *NSL-PN* to expand the network and download the *second layer* of protein interactions, thus obtaining the *Mito-CORE network* of the NSL complex. This is the protein interaction network built around the protein interactors of the NSL complex that are suggested to localise to the mitochondria.

The ***Mito-CORE network*** of the NSL complex was significantly enriched for the protein products of 180 PD associated genes, implicating a role for the NSL complex in the genetic risk of PD, via its mitochondrial functions. There are challenges to defining PD associated genes, here we have opted to include a panel of PD risk candidates from the recent GWAS ^15^ along with a set of diagnostic markers obtained from PanelApp v 1.68 (diagnostic grade genes for PD and Complex Parkinsonism) ^31^. Of course, caution must be taken in the interpretation of GWAS candidates, for which approaches to annotate causal genes at a given risk locus remain controversial ^39^. Considering these limitations, assessment of enrichment of a more stringent list of 15 genes, associated with Mendelian PD, has been carried out. It was observed that gene set enrichment is maintained (8/15 members represented) within the *Mito-CORE network*. Therefore, an enrichment of proteins encoded by PD associated genes within this network strongly reinforces the argument for a mitochondrial role of the NSL complex bridging sporadic and familial disease.

As a second approach, we have filtered the *first layer NSL-PN*, to retain the protein products of 12 PD associated genes present within the *first layer* of the *NSL-PN* (CSTA, PPP2R2B, SGF29, ATN1, VPS35, UBTF, SETD1A, CCAR2, CAMK2D, MAPT, CYLD and LRRK2). We used these 12 proteins to expand the network and download the *second layer*, thus obtaining the *PD-CORE network* of the NSL complex. This is the protein interaction network built around the protein interactors of the NSL complex that are coded by genes that present a genetic association with PD.

Functional enrichment analysis of the ***PD-CORE network*** showed the most significantly enriched biological processes are represented by the terms; *“mitochondrial cell death”, “mitochondrial protein localisation”* and the *“intracellular organisation of mitochondria*. Since PINK1 and Parkin are both absent from the refined *PD - CORE network*, the *PD – CORE network* has not been built upon the central components of the mitophagy system. This strengthens the link between dysregulation of mitochondrial dynamics in PD progression in sporadic and familial disease, with a low risk of ascertainment bias. As a final analysis, we have refined the *PD-CORE network*, enabling us to visualise and extract subnetworks that could provide mechanistic insight. We have first identified the entities within the *PD-CORE network* represented by the terms *“mitochondrial cell death”, “mitochondrial protein localisation”* and the *“intracellular organisation of mitochondria”*. We have then highlighted these within the network, using the Cytoscape (v.3.8.2) visualisation tool ^33^, to reveal the interactions with the NSL complex members that mediate this enrichment. The results of this final analysis point to OGT and WDR5 as possible key drivers of the PD risk associated with these mitochondrial processes.

Taken together, the findings of this work provide further evidence for a role of the NSL complex in driving PD pathogenesis. Specifically, we have illuminated that the PD association, at the protein level, is underpinned by a set of mitochondrial processes, strengthening a role for mitochondrial dynamics in both familial and sporadic disease. Finally, having isolated subnetworks relevant to this functional enrichment, we suggest OGT and WDR5 as candidates for further functional validation. This bioinformatics-led approach serves as a proof-of-principle, as an unbiased approach to extracting biologically meaningful information from genetic findings; to facilitate drug discovery.

## Acknowledgements

HPF and PAL were funded by Aligning Science Across Parkinson’s (grant number ASAP-000478) through the Michael J. Fox Foundation for Parkinson’s Research (MJFF). KK is funded by the Masonic Charitable Foundation. PAL and CM received funding from the Biomarkers Across Neurodegenerative Diseases Grant Program 2019, BAND3 (Michael J. Fox Foundation, Alzheimer’s Association, Alzheimer’s Research UK, and the Weston Brain Institute [grant number 18063]).

