## Supplementary Figures for "Protein network analysis links the NSL complex to Parkinson’s disease and mitochondrial biology"

### Slide 1
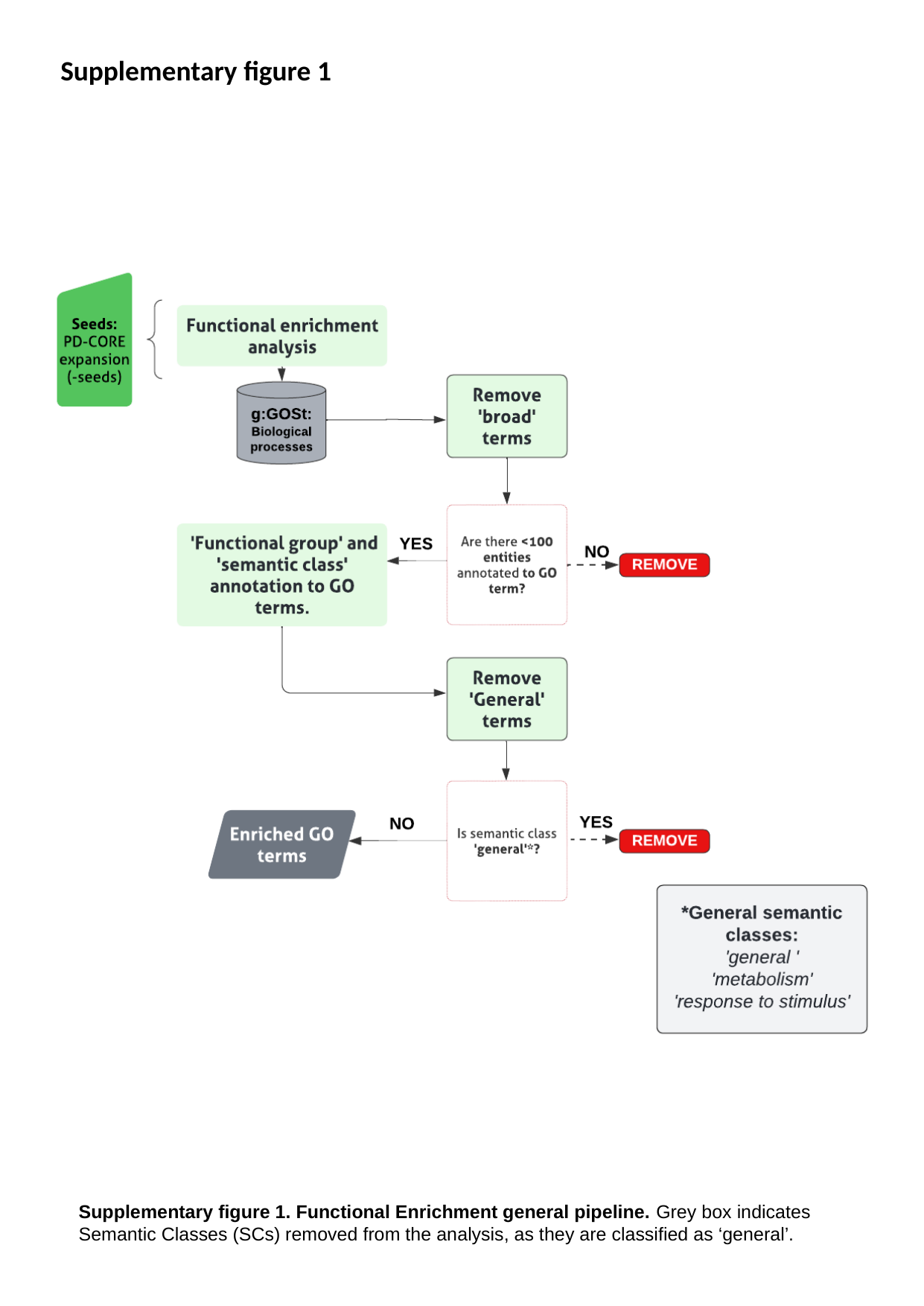

Supplementary figure 1
Supplementary figure 1. Functional Enrichment general pipeline. Grey box indicates Semantic Classes (SCs) removed from the analysis, as they are classified as ‘general’.

### Slide 2
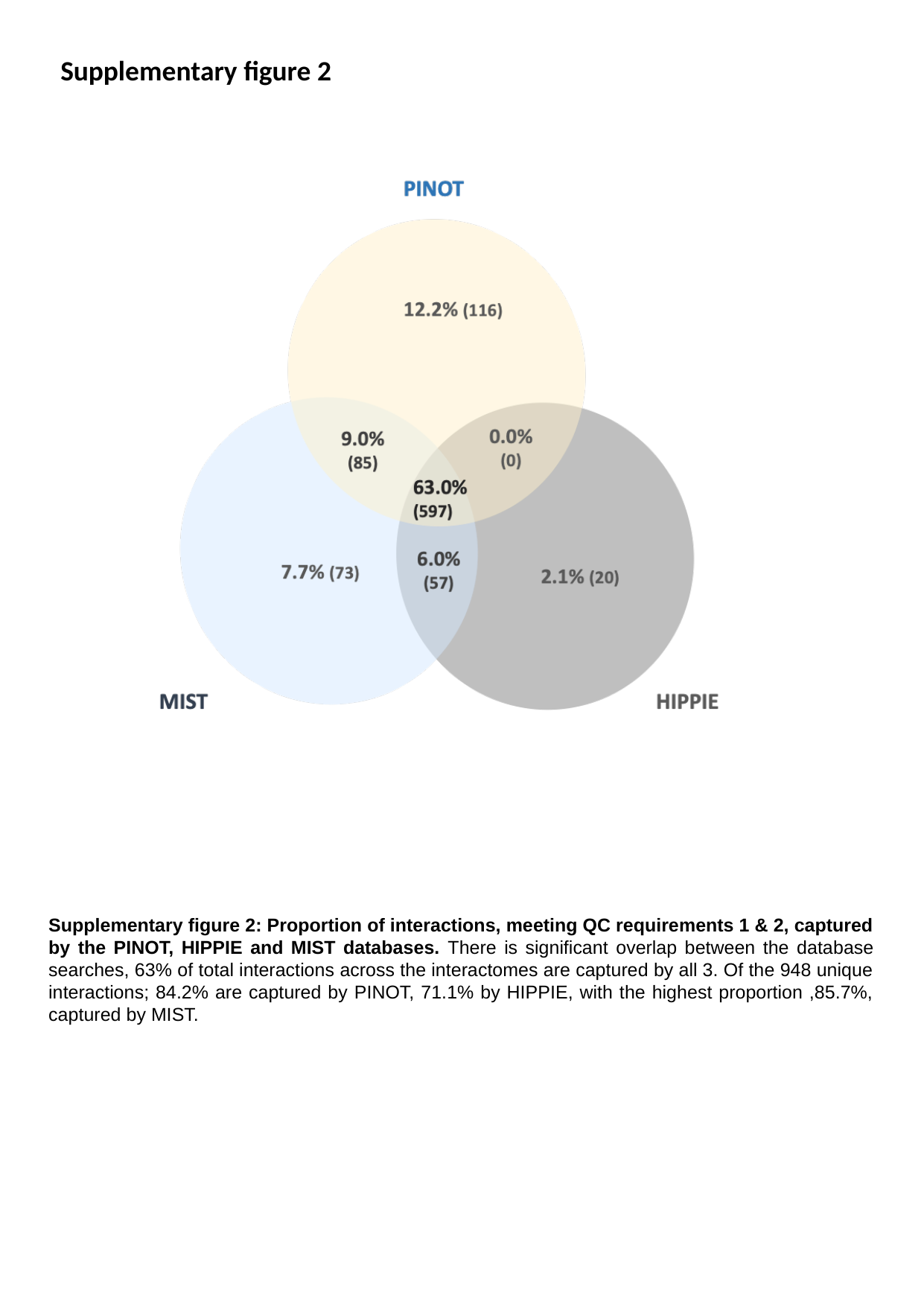

Supplementary figure 2
Supplementary figure 2: Proportion of interactions, meeting QC requirements 1 & 2, captured by the PINOT, HIPPIE and MIST databases. There is significant overlap between the database searches, 63% of total interactions across the interactomes are captured by all 3. Of the 948 unique interactions; 84.2% are captured by PINOT, 71.1% by HIPPIE, with the highest proportion ,85.7%, captured by MIST.

### Slide 3
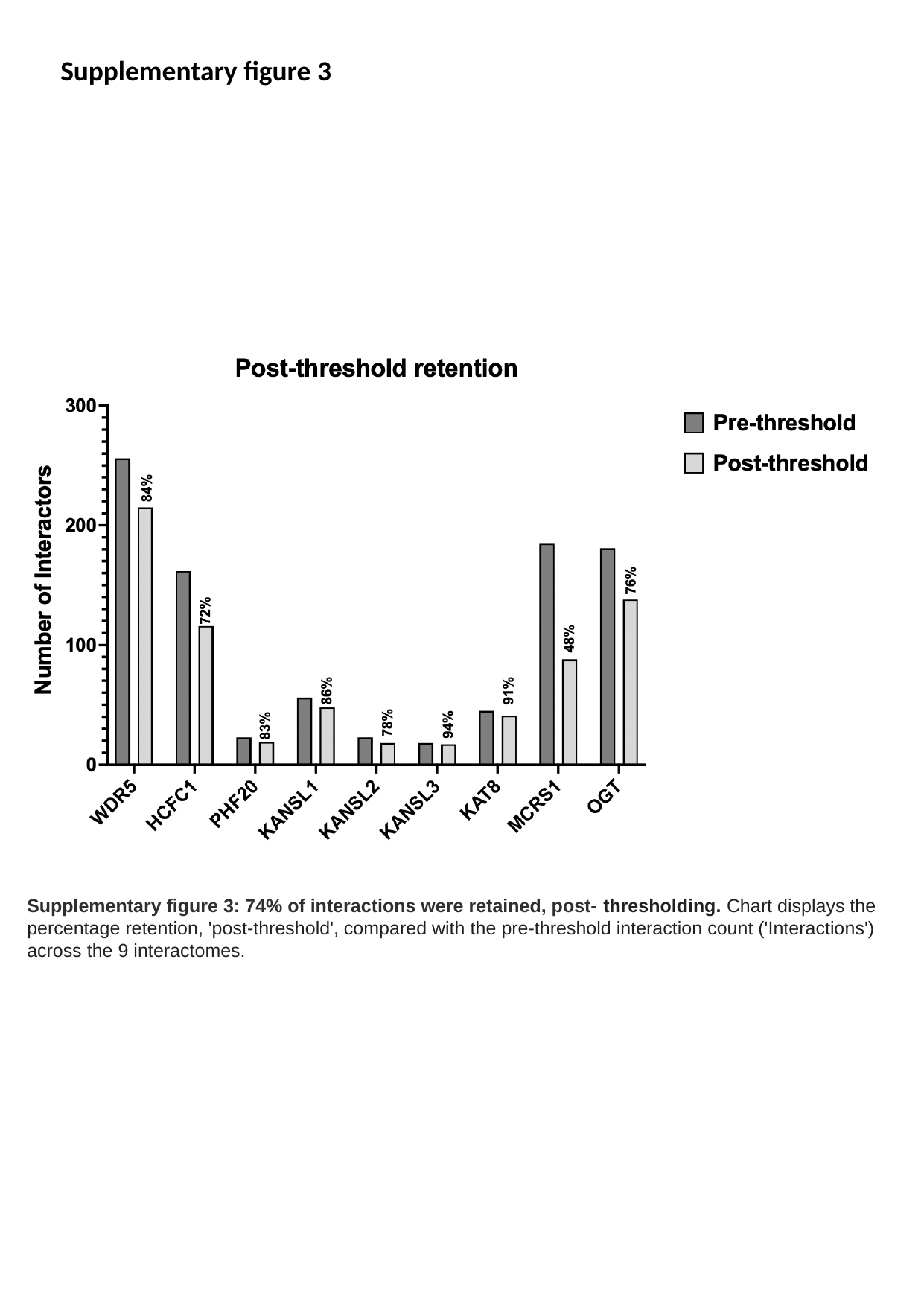

Supplementary figure 3
Supplementary figure 3: 74% of interactions were retained, post- thresholding. Chart displays the percentage retention, 'post-threshold', compared with the pre-threshold interaction count ('Interactions') across the 9 interactomes.

### Slide 4
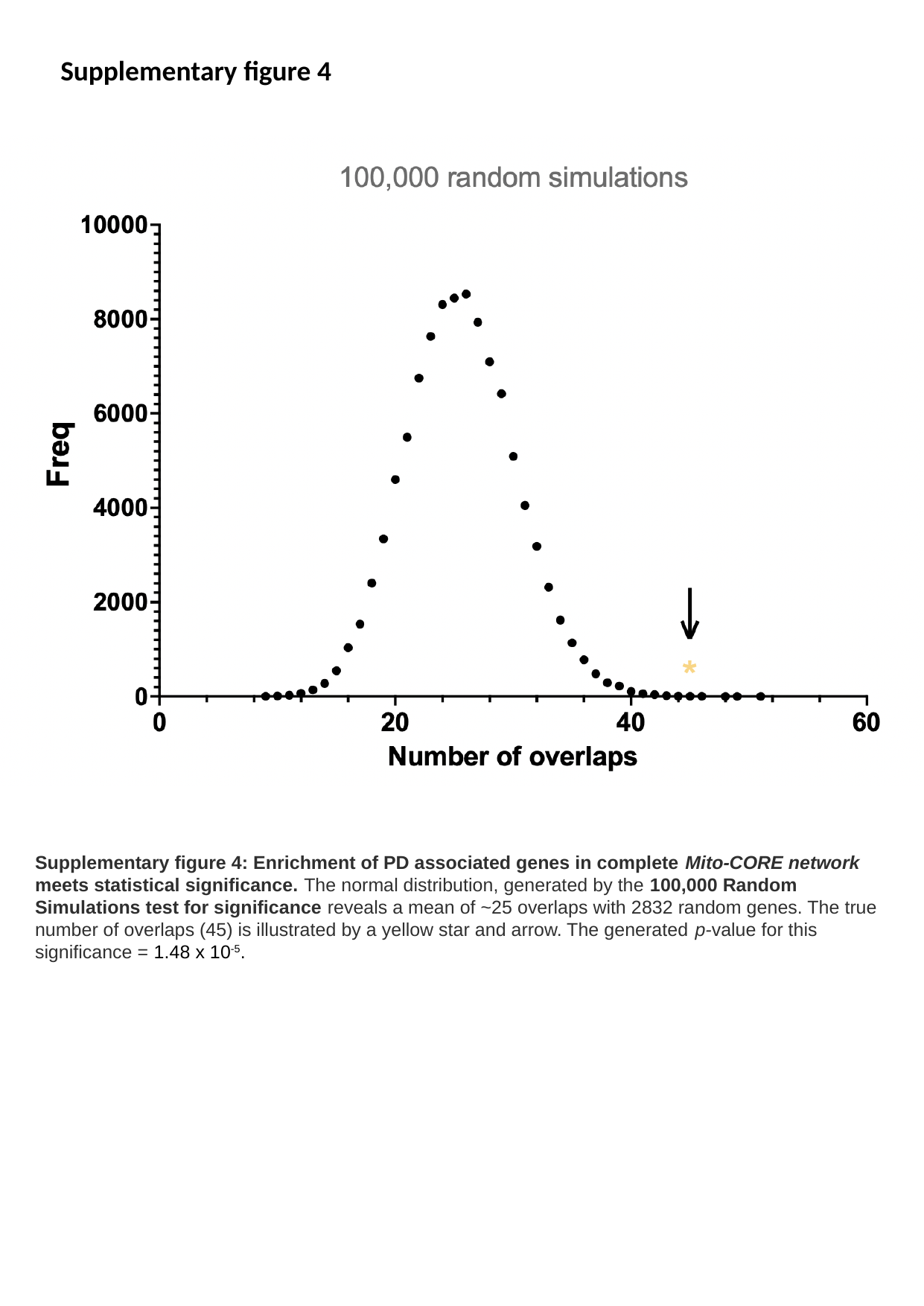

Supplementary figure 4
Supplementary figure 4: Enrichment of PD associated genes in complete Mito-CORE network meets statistical significance. The normal distribution, generated by the 100,000 Random Simulations test for significance reveals a mean of ~25 overlaps with 2832 random genes. The true number of overlaps (45) is illustrated by a yellow star and arrow. The generated p-value for this significance = 1.48 x 10^-5^.

### Slide 5
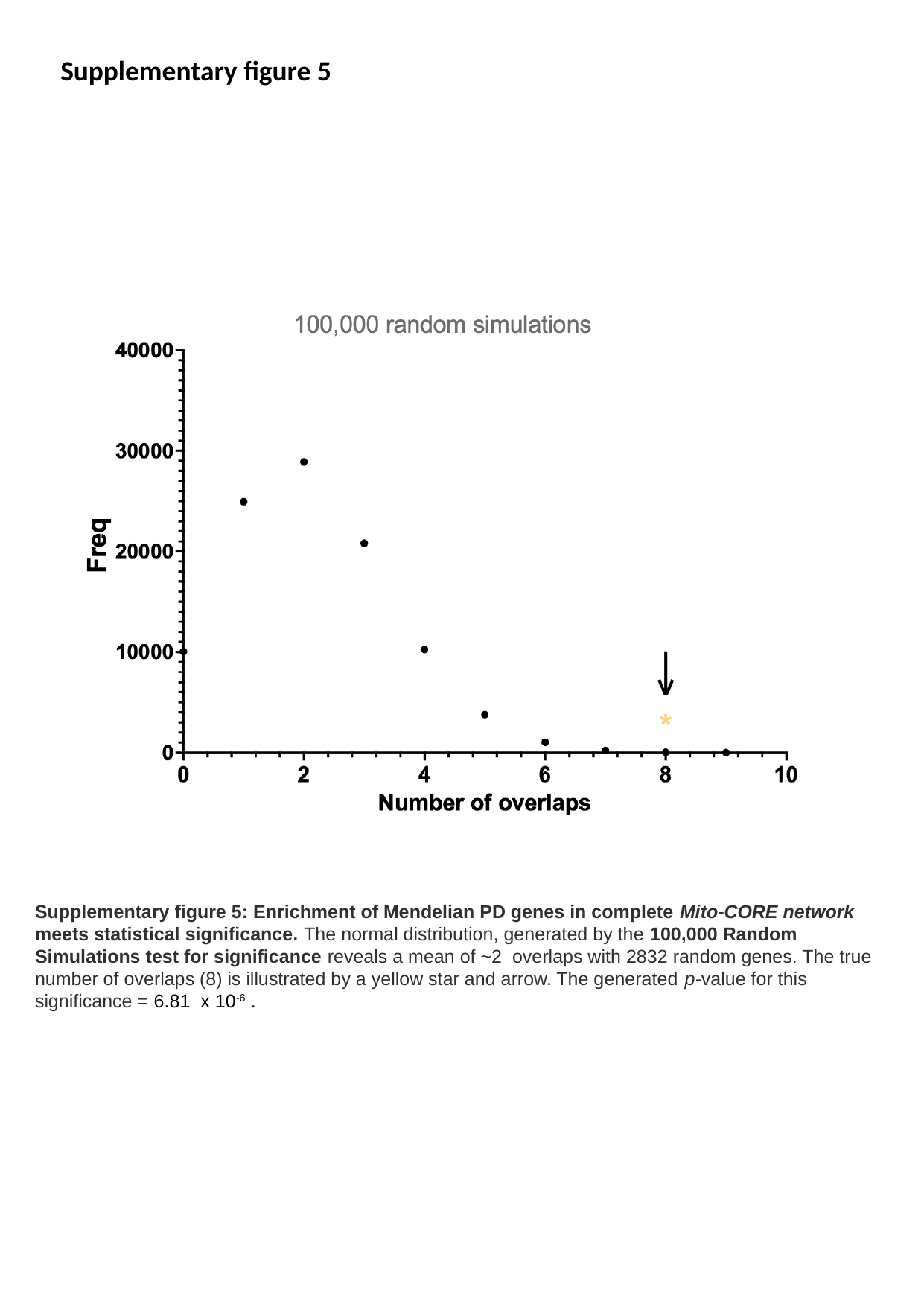

Supplementary figure 5
Supplementary figure 5: Enrichment of Mendelian PD genes in complete Mito-CORE network meets statistical significance. The normal distribution, generated by the 100,000 Random Simulations test for significance reveals a mean of ~2 overlaps with 2832 random genes. The true number of overlaps (8) is illustrated by a yellow star and arrow. The generated p-value for this significance = 6.81 x 10^-6^ .
